# Mining multi-omics data for molecular subtyping in gliomas facilitates precise clinical treatment

**DOI:** 10.1101/2023.05.25.542251

**Authors:** Peng Feng, Yawen Pan

## Abstract

Glioma is the most common malignancy in the skull, accounting for approximately 2% of adult systemic tumors. It is characterized by a high recurrence rate, high disability rate, and poor sensitivity to chemotherapy. Different subtypes of glioma exhibit varying prognosis and chemotherapy sensitivities. Currently, researchers extensively study the molecular classification of glioma based on transcriptomic features, facilitating the evaluation of prognosis. However, identification of tumor subtypes based on the single-layer-omics data have several limitations. In this regard, the present study aimed to identify new subtypes of glioma using three omics datasets for prognosis prediction. As a result, cluster A subtype was identified to be associated with lower activation of cell proliferation and better prognosis, while cluster B subtype exhibited higher infiltration of M2-type macrophages and higher activation of the epithelial-mesenchymal transition (EMT) pathway, potentially leading to a poorer prognosis. Furthermore, we employed the Weighted Gene Co-expression Network Analysis (WGCNA) algorithm and limma R package to identify driver genes for clusters A and B. Consequently, nine genes were identified as gene signatures for cluster A. Based on this finding, we quantified the two subtypes using the single-sample gene set enrichment analysis (ssGSEA) algorithm, where a higher score were linked to elevated tumor mutation burden (TMD) and signaling pathways related to cell proliferation. Low scores indicated enrichment of tumorigenesis-related pathways and poor prognosis. In silico drug screening suggested that AGI-6780 could be an effective compound for high cluster subtypes, based on brain tumor cell lines. Consequently, our study determined that the score can serve as an effective index to predict the prognosis of glioma.

## Introduction

Glioma, the most prevalent malignant brain tumor in adults, can be classified into five subtypes according to the 2016 WHO classification: IDH-mutant low-grade gliomas (LGGs) with chromosome 1p/19q codeletion, IDH-mutant LGGs without 1p/19q codeletion, IDH-mutant glioblastomas (GBMs), IDH wild-type LGGs, and IDH wild-type GBMs[1]. Intertumoral heterogeneity, based on the overall expression profile of tumors, has identified four subtypes of GBMs: anterior nerve (TCGA-PN), classical (TCGA-CL), mesenchymal (TCGA-MES), and neural (TCGA-NE)[2]. Despite advancements in multimodal therapy for gliomas, including surgery, radiotherapy, systemic therapy, and supportive therapy, the combination of adjuvant temozolomide and tumor treatment fields that uses low-intensity alternating electric fields remains an option. However, the overall prognosis for patients with glioma remains bleak, with a low long-term survival rate. Several factors, including advanced age, poor performance status, incomplete resection, and the absence of precise treatment strategies, have been identified as indicators of a poor prognosis[37]. Additionally, the associated morbidity, characterized by gradual deterioration in neurological function and quality of life, can cause immense hardship for patients, healthcare providers, and families[45]. Elderly patients, in particular, have a median survival of less than four months with only optimal supportive care[39]. The World Health Organization (WHO) updated its classification of central nervous system (CNS) tumors in 2021. This revision introduces significant changes that emphasize the role of molecular diagnostic methods in CNS tumor classification and highlight the importance of stratified reporting. This reporting style includes a comprehensive diagnosis, molecular information, CNS WHO classification, and tumor histopathological classification. Furthermore, the update incorporates several new tumor types identified through innovative diagnostic techniques like DNA methylome analysis[40]. In the case of gliomas, tumor classification serves as a valuable guide for precise treatment. Significant progress has been made in molecular characterization, aiding in tumor typing and facilitating precision treatment of gliomas. Mutations in isocitrate dehydrogenase 1 (IDH-1) and IDH-2, as well as O^6^-methylguanine DNA methyltransferase (MGMTp) methylation, have proven to be promising prognostic markers[41]. Treatment advancements have consequently increased the median survival rate to over 15 months in treated patients[43]. Therefore, given the promising efficacy displayed by numerous novel therapies for gliomas, it is crucial to identify new molecular subtypes that can lead to more sustainable treatment responses in patients suffering from this aggressive cancer, improving patient survival outcomes and extending overall survival, and enhancing patients’ quality of life.

Over the past five decades, the genetic approach to understanding cancer has dominated the research field. However, this perspective, focused solely on the genome, has limitations and tends to portray cancer as a highly heritable disease. Recent research suggests that cancer should be viewed as a disease in a multi-omics field and it is not heritable or completely heritable. It is now understood that the onset and development of cancer are affected by multiple factors, such as the exposome, metabolome, and genome. Notably, cancer-specific metabolism undergoes genetic alterations to support the growth and proliferation of cancer cells[32]. In this regard, relying solely on a single molecular dataset can only provide a partial understanding of the underlying causes of tumor growth. Multi-omics data have been utilized in some studies to classify patients with cancer and predict prognosis[33]. Consequently, the key to predicting tumor survival and recurrence prediction lies in performing comprehensive analysis of multi-omics data to reveal new characteristics of tumors. Multiple clustering algorithms designed for multi-data analysis enable the identification of novel molecular subtypes, facilitating the discovery or reconstruction of tissue-based tumor types. Molecular classifications that incorporate genetic and epigenetic alterations offer new insights into tumor treatment strategies and help to predict immunotherapy response. In the case of gliomas, multi-omics analysis can uncover previously undetected carcinogenic changes and guide the exploration of novel treatments. The utilization of the MOVICS[35] algorithm for multi-omics analysis offers promising prospects for prognostic evaluation and treatment of patients with glioma.

In this study, our primary objective is to investigate the relationship between tumor pathways, immune infiltration, and patient outcomes. Additionally, we aim to identify characteristic genes that define specific molecular subtypes and explore effective and less toxic therapeutic strategies. To achieve these goals, we performed multi-omics data clustering analysis to delineate distinct molecular subtypes. Consequently, nine gens were identified as cluster A-signed genes. Based on these genes, the ssGSEA [36] algorithm was employed to quantify different subtypes for prognosis of glioma. Notably, our findings indicate that this scoring system holds promise as an indicator for immunotherapy response and immune evasion. To comprehensively evaluate the efficacy of the scoring system, a thorough analysis of the immune microenvironment, copy number variation, and tumor mutation burden was conducted to compare the similarities and differences between the two subtypes. Our investigation encompassed multiple omics perspectives and involved external datasets for robust validation. Furthermore, we conducted in silico drug screening analyses using glioma cell lines to identify potential therapeutic targets specific to different prognosis subtypes. Furthermore, we performed computer-based drug screening analyses on glioma cell lines to identify potential therapeutic targets specific to different prognosis subtypes. In conclusion, we developed a robust scoring system that enables the identification of distinct tumor subtypes in gliomas. This scoring system exhibits predictive potential and offers valuable guidance for clinical decision-making regarding the treatment for glioma.

## Materials and methods

### Data sources

Transcriptome expression data and clinical data were collected from two sources: the China Glioma Genome Atlas (CGGA) database (http://www.cgga.org.cn) and the Cancer Genome Atlas (TCGA) database (https://portal.gdc.cancer.gov/)[21]. Following the integration of the available samples containing omics and clinical characteristics, 1,641 patients with glioma were enrolled in the subsequent analysis. Somatic mutations and clinicopathological features were obtained from the platform cBioPortal (https://www.cbioportal.org/). To validate our findings, we utilized external datasets including GSE184941, GSE4412, GSE7696, and CGGA301[29].

### Tumor infiltration of immune cells and tumorigenesis pathways

The CIBERSORT tool was utilized to assess the tumor microenvironment (TME)[3] and its relationship with immune infiltration. This approach enabled us to decode the intricate interplay between the samples and the matrix of immune cell invasion. Additionally, ssGSVA was performed to explore the associations between 50 tumor-related pathways and the samples.

### Multi-omics data clustering and subtype identification

In our analysis, we integrated data from 22 immune cells, 50 tumorigenesis pathways, and the gene expression profile of the sample to identify distinct molecular subtypes. To ensure the selection of an appropriate number of molecular subtypes, we performed evaluations using the cluster prediction index [4] and gap statistics [5]. Furthermore, ten separate cluster analyses were conducted employing various methods, including iClusterBayes, moCluster, CIMLR, IntNMF, ConsensusClustering, COCA, NEMO, PINSPlus, SNF, and LRA, to identify distinct molecular subtypes. Each algorithm had a similar silhouette score.

### Weighted gene co-expression network analysis (WGCNA)

The WGCNA was carried out to identify relevant modules associated with both clusters, [6]. The scale-free topology criterion was employed to calculate the soft threshold, and we determined the optimal value for it. The minimum module size was established as 30 genes. The Dynamic Tree Cut method was utilized to identify modules. The MEDissThres parameter was 0.25.

### Differentially expressed genes (DEGs) associated with molecular subtypes

Based on our multi-omics data analysis, the patients with glioma were classified into two distinct subtypes. The limma package was used to identify DEGs, with a significance level of P<0.01 and absolute fold changes greater than 1.

### Stratification of DEGs and construction of a rank index

The Boruta package was used for noise reduction and feature selection. Signature A was determined as genes exhibiting positive correlation with clustering (cor > 0.4) while Signature B comprised genes with negative correlation (cor < −0.4). Subsequently, the ssGSEA algorithm [7] was employed utilizing 9 genes from Signature A to construct a rank index quantifying the different subtypes.

### Analysis of Validation set and tumor microenvironment

The TIDE tool [8] was utilized to predict immunotherapy response among 1,692 patients with glioma based on TCGA data. Furthermore, we obtained transcriptome data for immune-related genes after PD-1 blockade from the IMvigor210 dataset [9]. Additionally, the score of the TME was evaluated using the estimate package (https://bioinformatics.mdanderson.org/estimate/index.html).

### Statistical analysis

Data analyses were carried out using the software R version 4.2.2 (http://www.r-project.org). For continuous data, the Wilcoxon test was done to measure differences between the two subtypes. The relationship between two variables was assessed using the Pearson correlation coefficient. Survival rates associated with different subtypes were compared using Kaplan-Meier survival analysis. A statistically significant difference was defined as P < 0.05.

## Results

### Two molecular subtypes were identified by MOVICS

Ten clustering algorithms-based MOVICS R package were used to identify molecular subtypes of glioma based on the multi-omics data (gene expression data, tumorigenesis-related pathways, and immune cells infiltration). These algorithms included iClusterBayes, moCluster, CIMLR, IntNMF, ConsensusClustering, COCA, NEMO, PINSPlus, SNF, and LRA. Composite physiologic index (CPI) and gender-age-physiology (GAP) scores indicated that there were two molecular subtypes, both of which obtained equal scores (Figure 1A, B, D). Figure 1C presents the distribution of various immune cell infiltrations. Cluster B exhibited significant enrichment in M2 macrophages, EMT, P53 pathways, and TGF_BETA pathways, which correlated with lower survival rates (Figure 1E). The DEGs of the two clusters were identified using the limma R package (Figure 1F).

**Figure 1.**
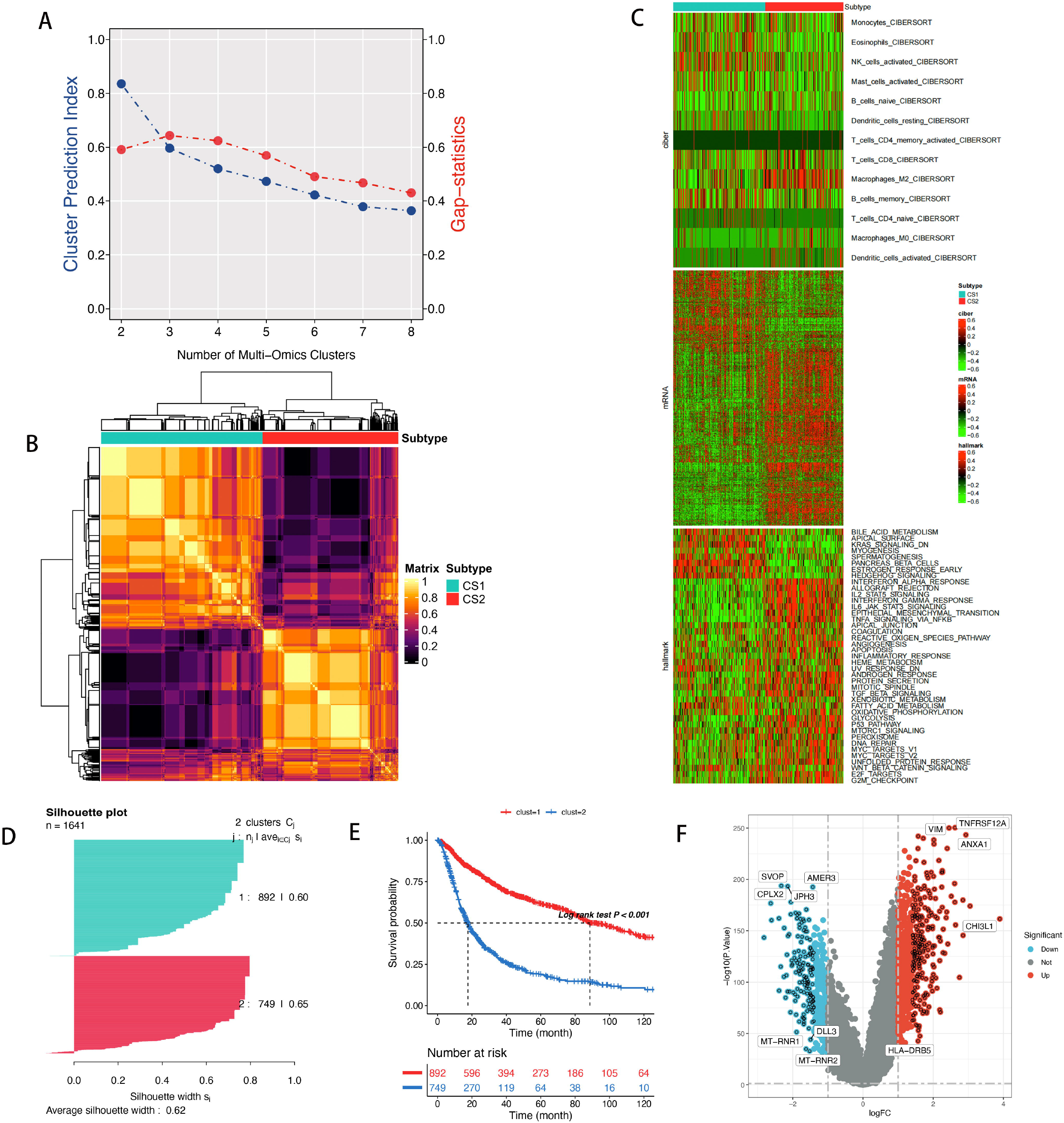
Identifying two molecule subtypes by using various clustering. (A) CPI analysis and gap-statistical analysis results. (B)Consensus matrix based on the various algorithms. (C)The landscape of molecular subtypes in the various immune cells infiltraion. (D) Silhouette-analysis evaluation. (E) The differential survival rate in the two clusters. (F) Differential expression genes in the two clusters.

### Transcription factor and copy number analysis

To figure out whether differences in the subtypes of glioma were underpinned by distinct molecular regulatory networks involving genetic and epigenetic regulators, we assessed eight glioma-specific transcription factors and 35 potentially regulated chromatin remodelers. According to Figure 2A, cluster A was likely to be regulated by HDAC5, SIRT5, FOXO3, KAT6B, KMT2A, OLIG2, SIRT1, and KAT2A, while Cluster B exhibited heightened activity of CARM1, FOXM1, KDM5C, PHF8, KDM4A, KDM5A, HDAC1, TGIF2, SIRT6, SIRT7 and GTF2B. The divergence in regulatory activity between these two clusters, which identifies the epigenetically driven transcriptional network, represents a crucial distinction. Copy number variation (CNA) stands as one of the prevalent large-scale genetic alterations found in cancer. Therefore, we proceeded to evaluate the copy number changes between the two clusters and found cluster B exhibited a significant increase in copy number compared to cluster A (Figure 2b). Moreover, the regions of notable copy number acquisition in cluster B were primarily identified as 7p11.2, 12q14.1, 1q32.1, and 4q12 region, as compared to clusters A. Chromosomal losses were concentrated in the 9p21.3 region (Figure 2C, D).

**Figure 2.**
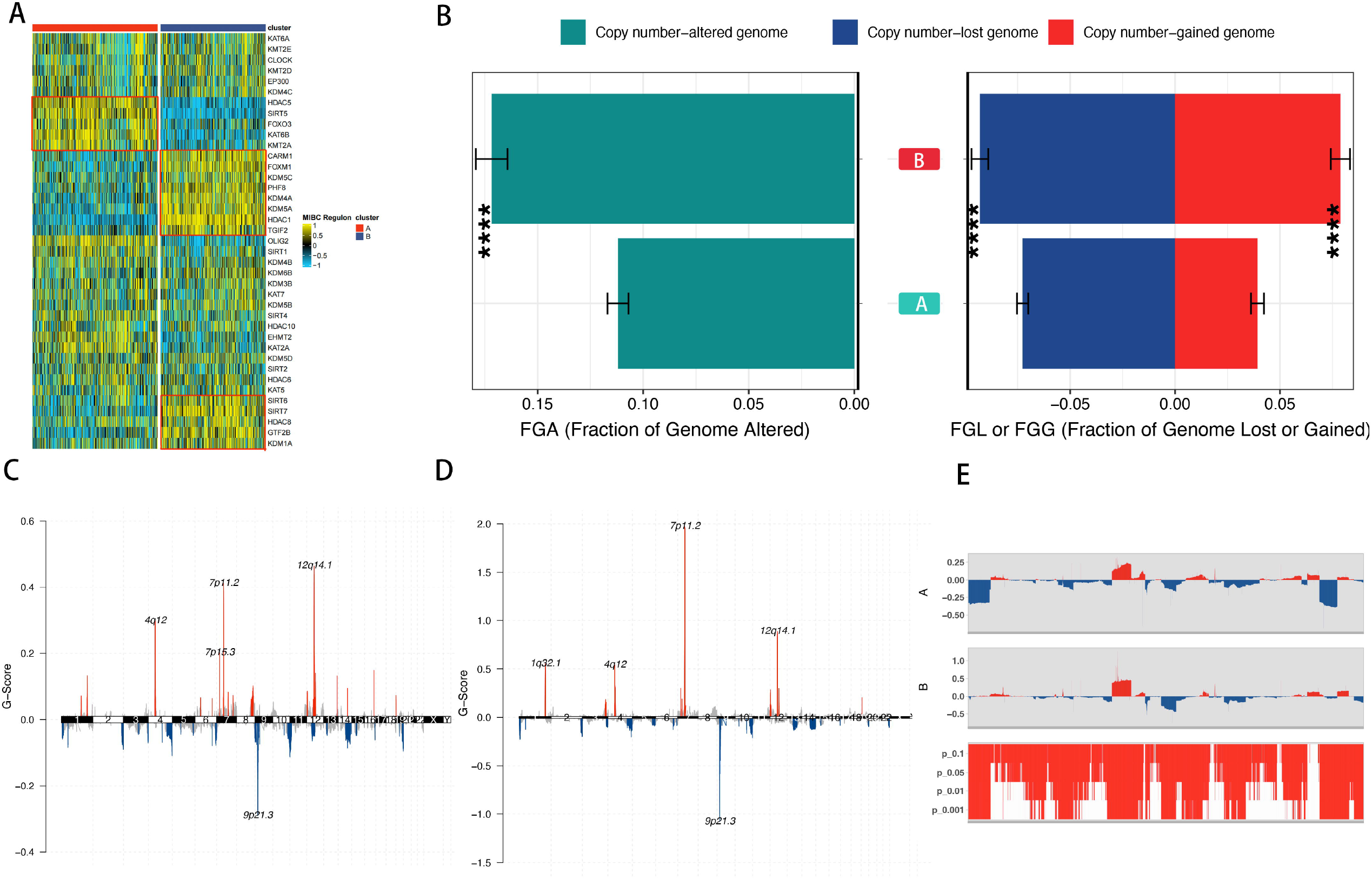
The alternation in the copy number and transcript factors activity. (A) Heatmap showing profiles of regulon activity for 6 gliomas-specific transcription factors and 33 chromatin remodeling potential regulons. (B) The distribution of fraction genome altered (FGA) and fraction genome gain/loss (FGA/FGG) in the to clusters. (C,D) The distribution of cluster A (C) and cluster B(D) in the copy number of genomic regions. (E) CNA plot shows the relative frequency of copy number gains (red) or deletions (blue) between the cluster A and cluster B of the gliomas cohort.

### WGCNA Network Module Mining

The scale-free topology criteria were used to calculate soft thresholds. The soft threshold power β = 5 (Figure 3A, B), Co-expression modules (cut height ≥ 0.25) were then identified based on phylogenetic trees (Figure 3C). Similar gene expression patterns were observed among modules belonging to the same branch when they underwent hierarchical clustering (Figure 3D). Consequently, these similar gene modules were combined, resulting in the formation of eight co-expression modules, denoted as brown, turquoise, black, blue, yellow, green, red, and gray (Figure 3E). Following this, the gene clusters were visualized, and inter-module correlations were analyzed (Figure 3F). Notably, the yellow module displayed a high correlation with cluster A, and thus it was identified as an important module for further investigation (Figure 3G) (Table 2).

**Figure 3.**
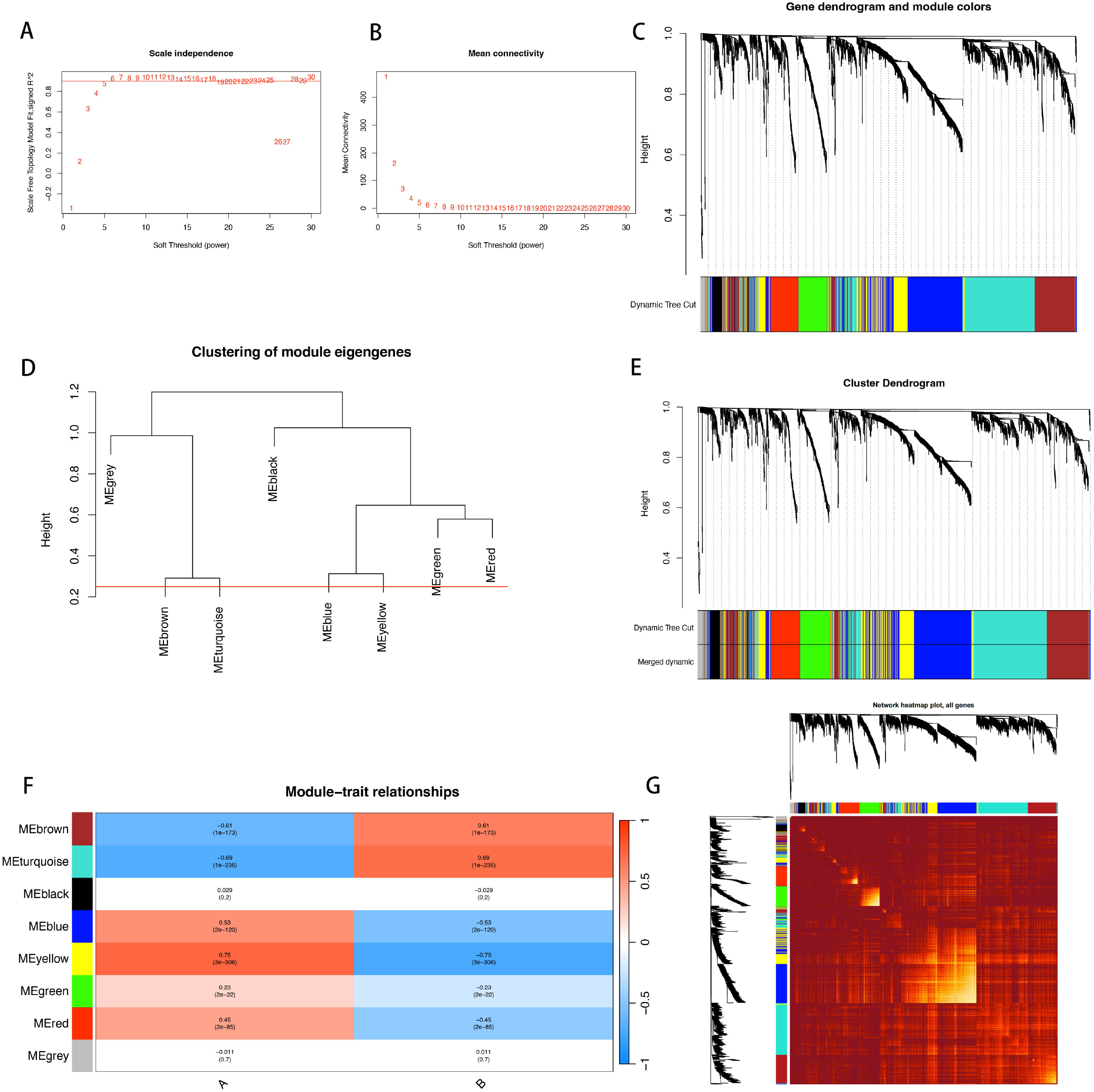
WGCNA analysis for the two clusters. (A) Confirming the best scale-free index for various soft-threshold powers (β). (B) The mean connectivity for various soft-threshold powers. (C) The gene tree map and nodule color. (D) Hierarchical clustering analysis. (E) The gene dendrogram is based on clustering. (F) Heatmap of the correlation between the module genes and the two clusters. (G) The heatmap of all genes.

### Dimensionality reduction and construction of characteristic genes

The correlation between the identified clusters and multi-omics data, including gene expression data, tumorigenesis-related pathways, and immune cell infiltration, was assessed using the Pearson correlation coefficient in this study. The analysis results demonstrated that Cluster B exhibited enrichment in cell proliferation-related pathways including EMT, P53 pathways, TGF_BETA pathways, and PI3K_AKT_MTOR_SIGNALING (Figure 4A). These results suggest that a strong activation of pathways related to cell proliferation could potentially be linked to unfavorable prognosis. Furthermore, the relationship between genes and clustering was investigated based on these differential genes and genes from the yellow module using the Pearson correlation coefficient. Consequently, nine genes were identified as A-signed genes (cor > 0.6, p < 0.01), while the signature B that did not meet these conditions (Figure 4B) (Table 3) was removed. Subsequently, the correlation between the two clusters of interest and clinicopathological factors was then analyzed.

**Figure 4.**
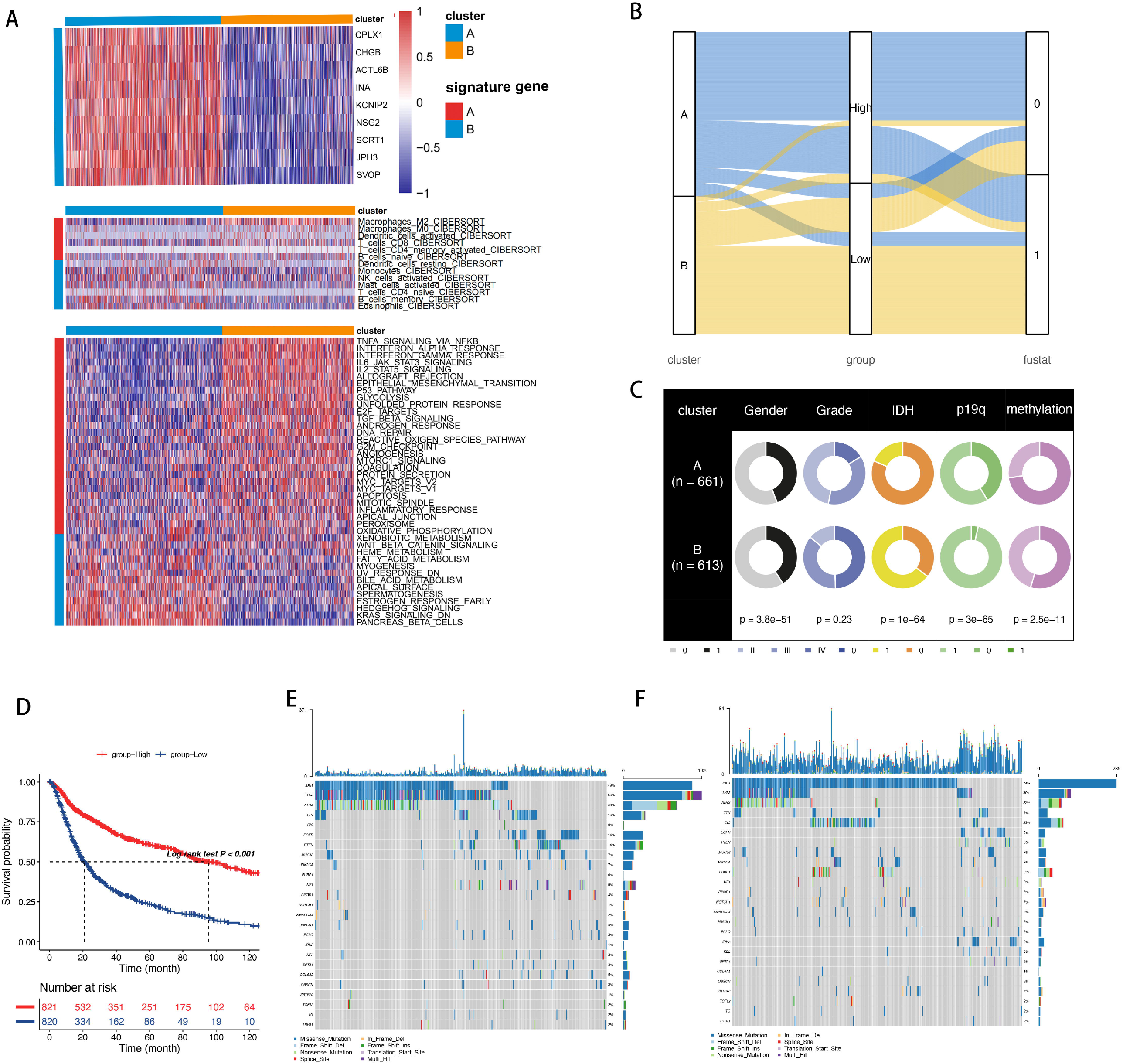
The alternation of genetic and mutation in the different subtypes. (A)Heatmap of 50 HALLMARKER pathways Immune cell infiltration and hub gene in the two clusters. (B)The distribution of clusters, and survival outcome in different score clusterss. (C) The distribution of clinicopathologic factors in the two clusters. (D)The differential survival rate in the different score clusterss. (G,H) The significant mutation genes in the high scores (E) and low scores(F).

The chi-square test results demonstrated that Cluster A had a low WHO grade with IDH1 mutation, 1p19q codeletion mutation, and less MGMT promoter (MGMTp) methylations mutation, indicating better survival outcomes compared with B. These findings are consistent with previous research. Furthermore, the relationship between clinical features and survival was evaluated using the TCGA and CCGA database, respectively, reaching a consistent conclusion (Supplementary Figure 3). These results suggest that our model serves as a reliable predictor of glioma prognosis and survival.

Based on the characteristics of Clusters A, the SSGSEA algorithm was utilized to calculate scores that could quantify the subtype. As shown in Figure 4B, the high clustering score predominantly overlapped Cluster A. We then evaluated survival in different scoring clusters and observed that the high-scoring clusters exhibited higher survival rates compared to the low-scoring clusters (Figure 4d). We also examined the expression differences of the nine signature genes in the A-B expression clusters, noting that the expression levels in Clusters A were generally higher than those in Clusters B (Supplementary Figure 1). Moreover, the receiver operation characteristic (ROC) curve values of all nine genes exceeded 0.8 (Supplementary Figure 2).

Next, we calculated the infiltration content of the nine core genes and their two sets across 22 immune cells and the enrichment of 50 pathways (Supplementary Figure 3A, B, D, E). The nine genes showed a strong correlation (Supplementary Figure 3C), and the high-scoring clusters exhibited strong anti-tumor efficacy, low enrichment of M2 macrophages, significant enrichment of B cells, monocytes, and CD4 memory T cells, as well as high immune activity. Conversely, the low-scoring clusters showed an abundance of M2 macrophages and significant enrichment in tumorigenesis-related pathways, especially signaling pathways such as TNFA_SIGNALING_VIA_NFKB, TGF_BETA_SIGNALING, EPITHELIAL_MESENCHYMAL_TRANSITION, and P53_PATHWAY, which may contribute to the poor prognosis of tumors.

### Analysis of tumor microenvironment and immune function

Building upon our previous analysis, we utilized our designated subtypes to examine disparities between the two subtypes in terms of the tumor environment and immune function (Figure 5A-C). Subtype B exhibited a greater infiltration of human leukocyte antigen-related genes (Figure7B), whereas subtype A displayed higher tumor purity than subtype B. Furthermore, we looked into the effects of the immune microenvironment on survival (Figure 7D-G). Consistent with our prior analysis (Figure 4C), subtype B exhibited more characteristics associated with high-grade tumors. Notably, progressive gliomas are known to have high interstitial and immune scores, indicating worse tumor purity. In our study, it was observed that higher tumor purity in gliomas was a potential key factor for poor survival, which aligns with the findings from previous research [19], In malignant tumors, excessive immune infiltration may cause immune storms and contribute to a poor prognosis. To gain further insights, we conducted Gene Ontology (GO) and KEGG pathways analysis (Figure 5H, I) on the clusters with high and low expression. These analyses highlighted several significant findings, including axon development, positive regulation of cell component biogenesis, cell growth, neuronal cell bodies, mitochondrial matrix, GTPase regulatory activity, and nucleoside-triphosphatase regulatory activity. The KEGG analysis indicated that the highly expressed clusters were involved in protein processing in the neurotrophic factor signaling pathway and endoplasmic reticulum, as well as enrichment in the MAPK signaling pathway. Additionally, GSVA enrichment analysis showed that EMTs were present in the top five pathways enriched in cluster B (Figure 5J). Conversely, cluster A exhibited enrichment in HALLMARK HEDGEHOG SIGNALING, and HALLMARK PANCREAS BETA CELLS (Figure 5K).

**Figure 5.**
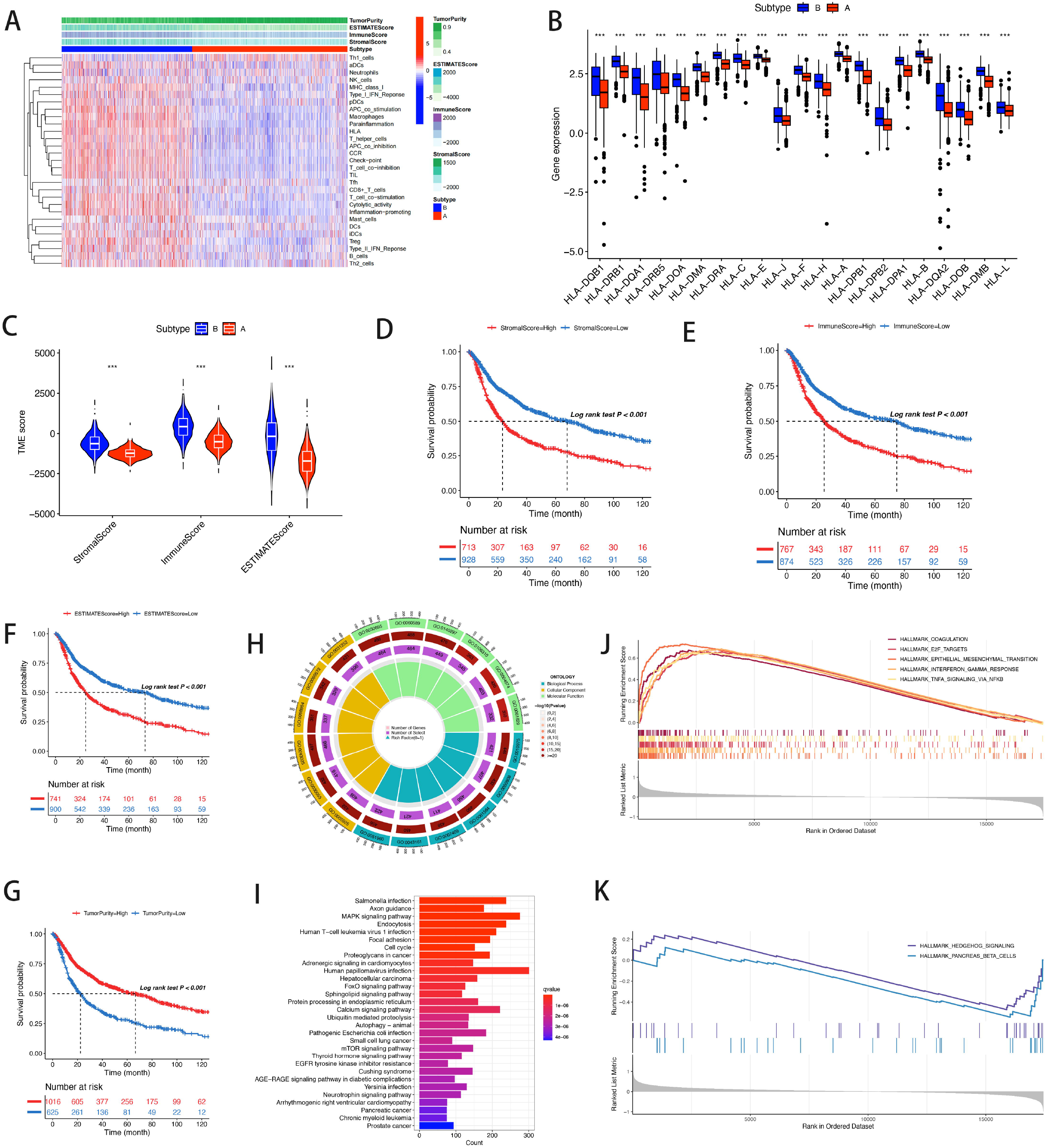
Analysis of immune microenvironment and function of different clusters. (A)Heatmap of tumor microenvironment and immune cell and function analysis between different clusters. (B,C) Differences between tumor microenvironment scoring(C) and HLA-associated gene expression(B) in different clusters. (D,E,F,G) The relationship between tumor microenvironment and survival. (H) GO analysis between different clusters. (I) KEGG analysis between different clusters. (J,K) The biological function of low(J) and high score clusterss(K).

### Tumor mutation burden (TMB) analysis

To acquire deeper understanding of the role of the model in glioma cell tumor mutation and immune evasion, we obtained mutational burden data for TCGA-GBM and TCGA-LGG and organized these data into an information matrix for analysis. Comparative analysis revealed that patients with glioma in the high clusters (Figure 4F) generally displayed a lower frequency of gene mutations than those in the low clusters (Figure 4E). However, there were exceptions to this trend. For example, genes such as IDH1, CIC, and FUBP1 exhibited higher mutation rates in the high clusters than in the lower clusters. Mutations in these genes may inhibit the progression of low-grade gliomas to high-grade gliomas and mediate the efficacy of targeted therapies. Furthermore, a differential analysis of TMB between patients in the high and low clusters was carried out. The results demonstrated a significant difference in TMB of glioma between these two groups, with patents in the high clusters exhibiting higher TMB than those in the low clusters (Figure 6A). Moreover, there was an inverse correlation (Spearman correlation = -0.237, P < 0.001) between the score and the degree of mutation (Figure 6B), indicating that as the score increased, the extent of mutation decreased. Additionally, we examined the prognostic implications of TMB. The survival time of patients in the clusters with high TMB differed significantly from that of patients in the clusters with low TMB, with patients in the former had significantly shorter survival time. Surprisingly, a positive correlation was observed between the score and patient survival outcome. These findings suggest that signature-related genes can mitigate tumor mutations in glioma cells, thereby improving patient prognosis and survival (Figure 6D, E).

**Figure 6.**
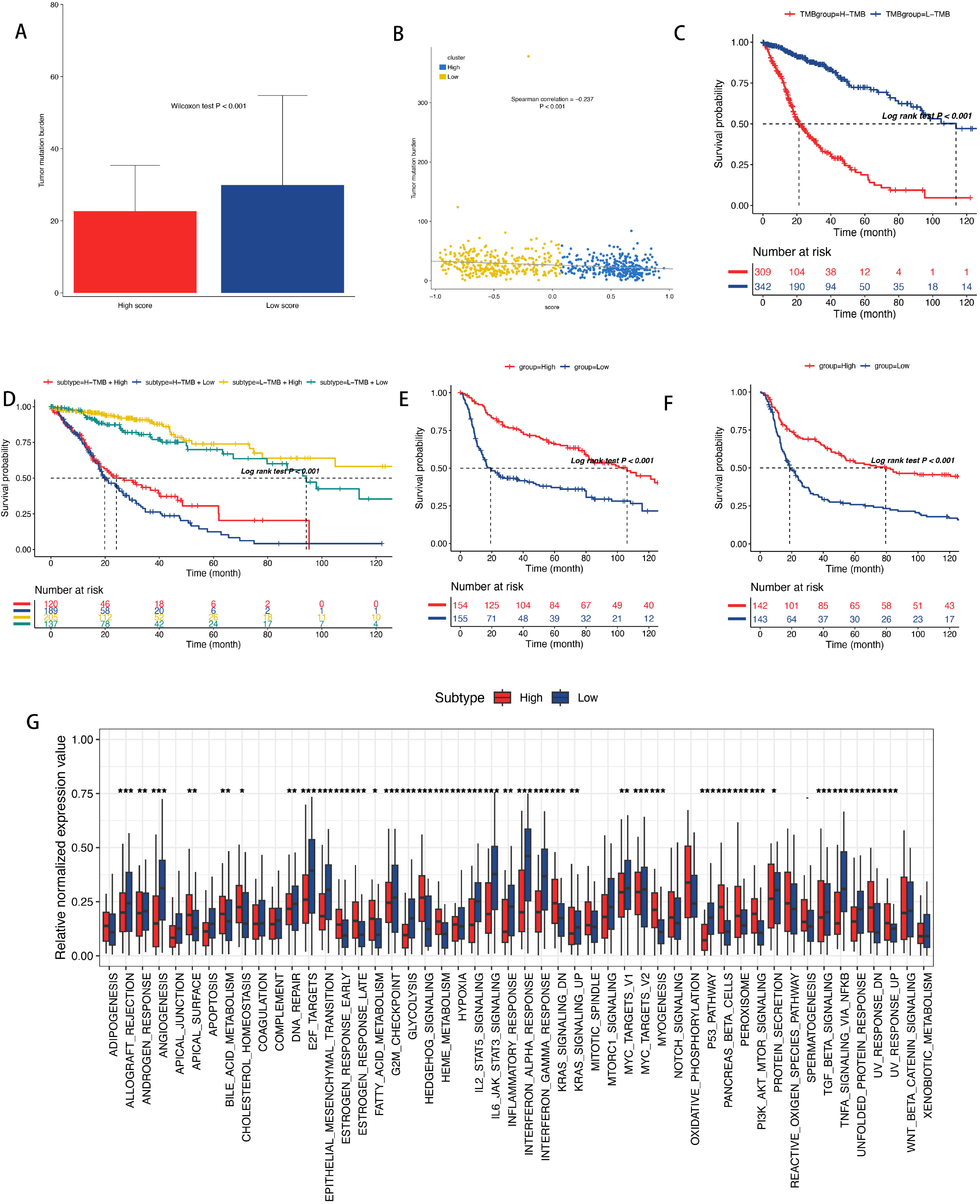
Tumor mutation burden analysis and subtype validation in the external cohorts.(A)Tumor mutation burden between high and low clusterss. (B)Correlation of score with tumor mutational burden. (C)The relationship between high and low mutation burden and survival. (D)The relationship between different mutation burden and survival in high and low clusterss. (E)Survival validation of external datasets(GSE184941&GSE4412&GSE7696(E) and CGGA301(F)). (G)Connections between external datasets and tumorigenesis pathways.

### Subtype validation in the external cohorts

To validate the subtypes in external cohorts, we obtained the external datasets (GSE184941, GSE4412, GSE7696 and CGGA301), combined them with the Gene Expression Omnibus (GEO) dataset, and performed survival validation using data from two different database sources. The ssGSEA was performed using 9 genes (referred to as signature A genes). Remarkably, in both cohorts, the high subclusters exhibited generally better prognoses than the low subclusters (Figure 6E, F). Surprisingly, in our validation set, an enrichment of EMT pathways was also found in the low subclusters, which aligns with the findings from our initial cohort.

### The value of in-silicon drug screening and scoring in immunotherapy response prediction

To explore more effective treatment strategies for the high and low clusters, the GDSC data was utilized to identify compounds that were sensitive to subtypes with different scores. Based on the 9 A-signature genes, the ssGSEA was conducted to score glioma cell lines. Interestingly, these 9 A-marker genes displayed significant expression at the transcriptional level of glioma cell lines (Figure 7A, B). Through susceptibility analysis, we observed in sensitivity to PHA-793887 in the high-scoring clusters, while the low-scoring clusters showed greater benefit from the potent selective IDH2 R140Q mutant inhibitor AGI-6780 (Figure 7C, D) (Table 4). Considering the growing application of immunotherapy in tumor treatment and its significant impact on cancer patient survival, particularly the use of PD-1 or PD-L1-specific monoclonal antibodies, we utilized the TIDE website[8] to predict the response to cancer immunotherapy among 1,691 patients from the TCGA database. Based on the predicted results, the high-scoring clusters exhibited higher objective response rate of TCGA anti-PD-L1 therapy (55.2%) compared to the low clusters (41.2%) (Figure 7E, F).

**Figure 7.**
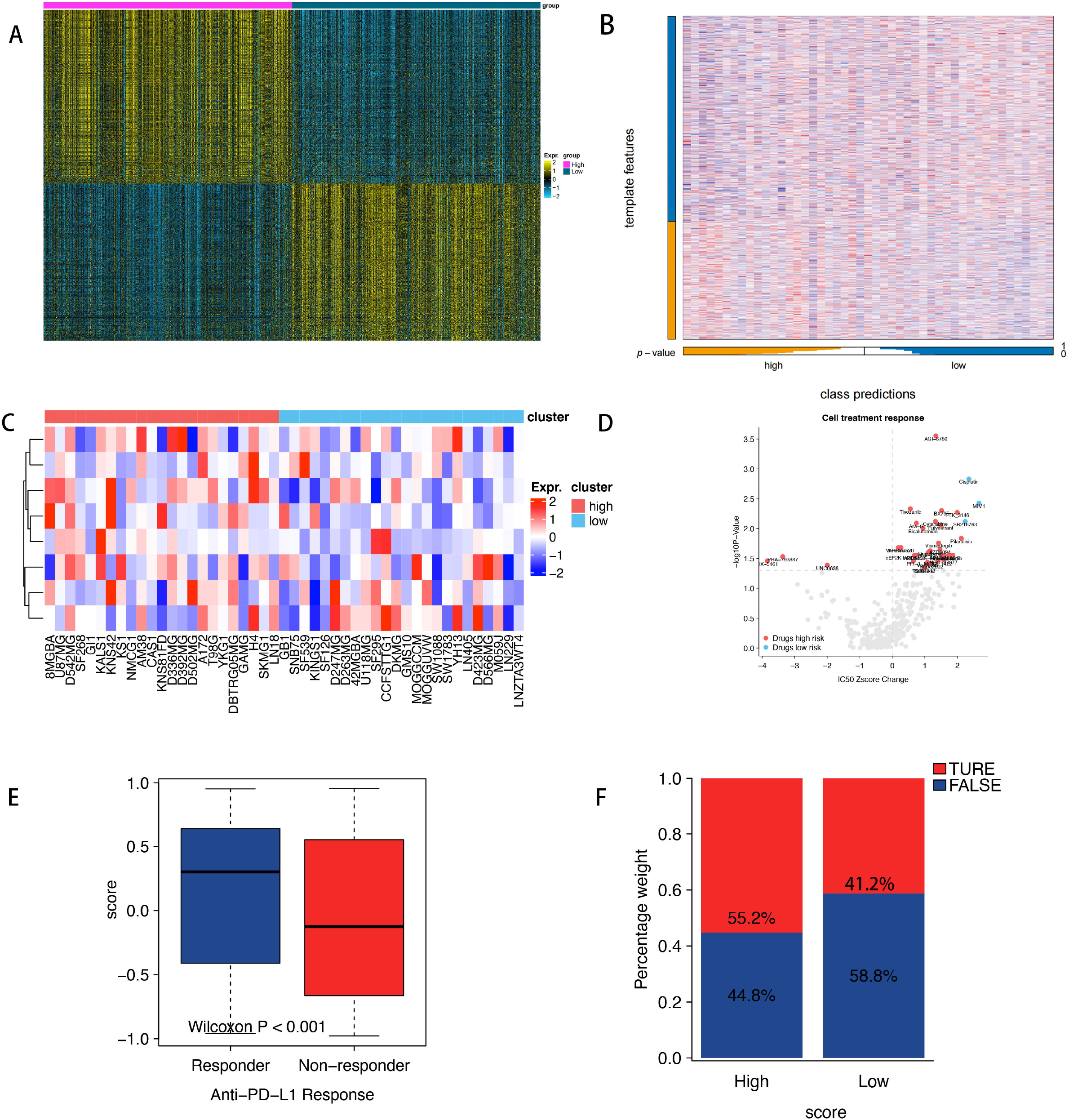
Drug screening and immunotherapy prediction between different clusters. (A)Heatmap of subtype-specific upregulated biomarkers using limma for identifying the two clusters. (B) Heatmap of NTP. (C) Heatmap shows the expression pattern of 67 genes in cancer cell lines showing low or high score clusters. (D)The figure summarizes the relative changes of the ic50z score and P values in selected gliomas cell systems at low or high score clusters which can be treated using the designated compounds based on the cancer drug susceptibility genomics (gdsc) database. The red spot clearly shows the high sensitivity drugs of high score clusters and the blue point clearly shows the high sensitivity drugs of low score clusters. (E) The distribution of score in the different anti-PD-1 response. (F) The response proportion of anti-PD-L1 immunotherapy (response/Non-response and stable disease (SD)/progressive disease (PD)) in the different score clusterss in the TIDE website after the prediction.

## Discussion

Brain cancer, particularly glioblastoma, is a highly malignant and aggressive form of cancer. Despite advancements in treatment, such as combination surgery, radiotherapy, and temozolomide therapy [20], the prognosis for glioblastoma remains poor, leading to high mortality rates. Additional therapies, including molecular targeted therapy and immunotherapy, have been explored for treating glioblastoma, but their effectiveness is limited. Molecular analysis has proven valuable in classifying brain cancer into distinct multiple molecular subtypes, providing more accurate prognostic predictions and treatment guidance. Comprehensive molecular classifications based on gene expression profiles at the transcriptome level have significantly improved our understanding of brain cancer biology. Malignant transformation involves multiple molecular alterations at various levels. To unravel the heterogeneity of brain cancer, we conducted clustering analysis using multiple datasets to reveal diverse molecular subtypes. In our study, the application of ten clustering methods identified two molecular subtypes characterized by distinct prognosis and activity in tumorigenesis-related pathways. Specifically, cluster A showed a better prognosis, while cluster B was significantly enriched in M2 macrophages and EMT, P53 pathways, and TGF_BETA pathways, resulting in poorer prognosis. Previous studies have consistently demonstrated the association between different subtypes and patient survival. In line with these findings, our study revealed higher mutation rates of IDH1, CIC, and FUBP1 in the high-expression clusters compared to the low-expression clusters. Drug screening further indicated that high-expression clusters benefited from potent selective inhibitors targeting IDH, whereas low-expression clusters demonstrated lower responsiveness, which may be related to mutations in IDH1. It has been reported that the N-terminal region of the IDH1 protein interacts with ATXN1[10] to form a transcriptional inhibitory complex. In vitro studies have suggested that the expansion of polyglutamine in ATXN1 may alter the inhibitory activity of this complex. Mutations in CIC and FUBP1 are associated with oligodendroglioma[11][12][13]. The lower expression of ATXN family genes in the low-expression clusters of high-resolution expression clusters reduces their interaction with CIC, thereby inhibiting the occurrence of oligodendroglioma.

Our study showed that cluster A has a good prognosis for glioma patients, which can be attributed the low activity of tumorigenesis-related pathways and low infiltration of M2 macrophage, indicating potential effectiveness in immunotherapy. Conversely, cluster B showed worse prognosis, which may be related to the significant enrichment of M2 macrophages and EMT, P53 pathways, and TGF_BETA pathways. It may also be affected by the activation of cell proliferation pathways regulated by CARM1, FOXM1, KDM5C, PHF8, KDM4A, KDM5A, HDAC1, TGIF2, SIRT6, SIRT7, and

GTF2B. Our assessment of copy number alterations (CNAs) between the two clusters revealed significant copy number loss primarily concentrated in the 9p21.3 region within cluster B. Loss of 9p21.3 not only fails to effectively block tumor cell proliferation but also facilitates immune evasion while disrupting cell-intrinsic and cell-extrinsic tumor suppressor programs.

As previously reported, the loss of copy number in genes associated with the MAPK pathway and MHC pathway can be used as a potential indicator for predicting the response to immunotherapy. Notably, the EGFR gene is essential to the regulation of PD-L1 expression[14][15]. Identifying tumors with high EGFR levels and increased lymphocyte infiltration could serve as a valuable approach to identifying individuals with cytotoxic immunophenotypes, who are more likely to respond favorably to PD-1 therapy. In this study, a comparison of gene CNAs between the two clusters revealed a significant loss of INF family genes located in the 9p21.3 region in cluster B compared with cluster A (TableS1). This observation suggests that genomic alterations impacting the copy number of the INF family genes [16][17][18] might enable cluster B to evade immune surveillance and promote tumorigenesis.

WGCNA was then performed and the limma R package was used to identify modular genes and DEGs between the two clusters. Based on the gene characteristics related to the enrichment of tumorigenesis pathways and immune cell infiltration, we performed ssGSEA to quantify ICI clusters.

Subsequently, patients were split into high- and low-score subgroups based on the median scores obtained. Analysis demonstrated that the high-score group exhibited a significant overlap with cluster A, while the low-score group showed a stronger association with cluster B, which is consistent with our initial hypothesis. Our study demonstrated that high-scoring patients exhibited elevated expression of immunoactivity-related features, while low-scoring patients displayed pronounced infiltration of M2-type macrophages and activation of EMT, P53 pathway, and TGF_BETA pathways. The distinguishing features between these two clusters exhibited significant differences and correlated with the rate of responses to immunotherapy. To validate the accuracy of our prediction model in determining survival distributions within high-scoring and low-scoring groups, we utilized two external cohorts from databases GEO and CCGA.

Gliomas, known for their high malignancy, currently rely on surgical resection combined with chemotherapy and radiation therapy as the primary treatment approach. However, these conventional treatments often lead to severe side effects, including potentially fatal outcomes, and their efficacy remains limited. Therefore, there is an urgent need for the creation of new and efficient strategies that have minimal negative impacts. Cancer immunotherapy has emerged as a promising avenue to either treat cancer or improve its prognosis. Various immunotherapy medications have been tested for gliomas, demonstrating potential therapeutic advantages with fewer side effects compared to traditional chemotherapy and radiation therapy. We determined that the low clustering subtype exhibited the most favorable potential for immunotherapy. To support this, TCGA cohorts were used and their correlation with scores were evaluated. The results revealed that the high-score subgroup displayed a higher likelihood of response to anti-PD-L1 treatment compared to the low-score subgroup in both cohorts. Furthermore, the subplot analysis indicated a high probability of immunotherapy response in the high clusters. Additionally, drug screening predicted that the selective IDH2 R140Q mutant inhibitor AGI-6780 exhibited potent properties in the low clusters. Our findings suggest that low clustering may benefit from the use of potent selective IDH2 R140Q mutant inhibitors.

We established a new score-based signature for predicting glioma based on multi-omics data and the collective outcomes from ten algorithms. High-score cluster which was characterized by the activation of inflammation-related pathways and increased expression of immune checkpoint proteins was associated with improved survival rates. Moreover, the high-score cluster displayed a higher probability of responsiveness to immunotherapy. The model in the present study can effectively predict the efficacy and prognosis of anti-PD-L1 treatment. However, the validation of this scoring system necessitates prospective analysis based on a sizable group of patients with glioma, and the effectiveness of anti-PD-L1 treatment for high-score cluster should be validated through appropriate preclinical models and future clinical trials. In summary, the findings of this study shed light on the intricate interplay between genetic and epigenetic events in tumor immunophenotyping for glioma. the implementation of multi-omics analysis for gliomas can help to offer patients more accurate clinical treatment options and improved prognosis prediction.

## Supporting information

**Supplementary Figure:**

**Figure S1:** The expression of core genes in different clusters.

**Figure S2:** Roc correlation map of core genes.

**Figure S3:** Differential heat map of core genes in tumorigenesis pathways and immune cell infiltration in different clusters (A)Differences in 22 types of immune cells in different clusters. (B)Correlation heat map of core genes with 22 immune cells. (C)Core gene-related heat map. (D)Differences between different clusters in 50 tumorigenesis pathways. (E)Correlation heat map of core genes with 50 tumorigenesis pathways.

**Figure S4:** The expression of core genes is the relationship between survival and survival.

**Figure S5:** Clinical features and survival of different clusters.

**Supplementary Table:**

**Table S1:** List of genetic mutations.

**Table S2:** List of yellow module genes.

**Table S3:** List of intersections of differential genes with core genes.

**Table S4:** List of drug predictions.

## References

[1] Brat DJ, & Pachter L. (2015). Comprehensive, Integrative Genomic Analysis of Diffuse Lower-Grade Gliomas.

[2] Verhaak, R. G. W., Hoadley, K. A., Purdom, E., Wang, V., Qi, Y., & Wilkerson, M. D., et al. (2010). Integrated genomic analysis identifies clinically relevant subtypes of glioblastoma characterized by abnormalities in pdgfra, idh1, egfr, and nf1. Cancer Cell, 17(1), 98–110.

[3] Chen B, Khodadoust MS, Liu CL, Newman AM, Alizadeh AA. Profiling Tumor Infiltrating Immune Cells with CIBERSORT. Methods Mol Biol. 2018;1711:243–259. doi:10.1007/978-1-4939-7493-1_12.

[4] Chalise P., Fridley B.L. Integrative clustering of multi-level omic data based on non-negative matrix factorization algorithm. PLoS ONE. 2017;12:e0176278. doi: 10.1371/journal.pone.0176278.

[5] Hastie T., Tibshirani R., Walther G. Estimating the number of data clusters via the gap statistic. J. R. Stat. Soc. B. 2001;63:411–423.CrossRefGoogle Scholar

[6] Langfelder P, Horvath S. WGCNA: an R package for weighted correlation network analysis. BMC Bioinformatics. 2008;9:559. Published 2008 Dec 29. doi:10.1186/1471-2105-9-559

[7] Hänzelmann S, Castelo R, Guinney J (2013). “GSVA: gene set variation analysis for microarray and RNA-Seq data.” BMC Bioinformatics, 14, 7. doi: 10.1186/1471-2105-14-7.

[8] Jiang, P., Gu, S., Pan, D., Fu, J., Sahu, A., Hu, X., et al. (2018). Signatures of T Cell Dysfunction and Exclusion Predict Cancer Immunotherapy Response. Nat. Med. 24, 1550–1558. doi:10.1038/s41591-018-0136-1.

[9] Mariathasan S, Turley SJ, Nickles D, et al. TGFβ attenuates tumour response to PD-L1 blockade by contributing to exclusion of T cells. Nature. 2018;554(7693):544–548. doi:10.1038/nature25501.

[10] Oncotarget, ALBANY, v., n., & supl. (2012). Frequent atrx, cic, fubp1 and idh1 mutations refine the classification of malignant gliomas. Oncotarget, 3(7), 709–722.

[11] (2011). Mutations in cic and fubp1 contribute to human oligodendroglioma. Science, 333(6048), 1453–1455.

[12] Yang, R., Chen, L. H., Hansen, L. J., Carpenter, A. B., Moure, C. J., & Liu, H., et al. (2017). Cic loss promotes gliomagenesis via aberrant neural stem cell proliferation and differentiation. Cancer Research, 6097.

[13] Oncotarget, ALBANY, v., n., & supl. (2012). Frequent atrx, cic, fubp1 and idh1 mutations refine the classification of malignant gliomas. Oncotarget, 3(7), 709–722.

[14] Proteogenomics of diffuse gliomas reveal molecular subtypes associated with specific therapeutic targets and immune-evasion mechanisms. Nature Communications.

[15] Erel-Akbaba, G., Carvalho, L. A., Tian, T., Zinter, M., Akbaba, H., & Obeid, P. J., et al. (2019). Radiation-induced targeted nanoparticle-based gene delivery for brain tumor therapy. ACS Nano.

[16] Gao, J., Shi, L. Z., Zhao, H., Chen, J., Xiong, L., & He, Q., et al. (2016). Loss of ifn-γ pathway genes in tumor cells as a mechanism of resistance to anti-ctla-4 therapy. Cell.

[17] Zaretsky, J. M., Garcia-Diaz, A., Shin, D. S., Escuin-Ordinas, H., Hugo, W., & Hu-Lieskovan, S., et al. (2016). Mutations associated with acquired resistance to pd-1 blockade in melanoma. New England Journal of Medicine, 819.

[18] Shin, D. S., Zaretsky, J. M., Escuin-Ordinas, H., Garcia-Diaz, A., Hu-Lieskovan, S., & Kalbasi, A., et al. (2016). Primary resistance to pd-1 blockade mediated by jak1/2 mutations. Cancer Discovery, 188.

[19] Zhang, C. B., Cheng, W., Ren, X., Wang, Z., Liu, X., & Li, G., et al. (2017). Tumor purity as an underlying key factor in glioma. Clinical Cancer Research, clincanres.2598.2016.

[20] Phimister E G, Das S, Marsden P A. Angiogenesis in Glioblastoma[J]. New England Journal of Medicine, 2013, 369(16):1561–1563.

[21] Zhao, Z., Zhang, KN., Wang, QW., et al. Chinese Glioma Genome Atlas (CGGA): A Comprehensive Resource with Functional Genomic Data from Chinese Glioma Patients (2021). Genomics, Proteomics & Bioinformatics 19(1):1–12.

[22] Zhao, Z., Meng, F., Wang, W., et al. (2017). Comprehensive RNA-seq transcriptomic profiling in the malignant progression of gliomas. Scientific data 4, 170024

[23] Bao, Z.S., Chen, H.M., Yang, M.Y., et al. (2014). RNA-seq of 272 gliomas revealed a novel, recurrent PTPRZ1-MET fusion transcript in secondary glioblastomas. Genome research 24, 1765–1773

[24] Zhang, K., Liu, X., Li, G,. Clinical management and survival outcomes of patients with different molecular subtypes of diffuse gliomas in China (2011– 2017): a multicenter retrospective study from CGGA. Cancer Biol Med 19, 1460-1476. 10.20892/j.issn.2095-3941.2022.0469.

[25] Zhao, Z., Zhang, KN., Wang, QW., et al. Chinese Glioma Genome Atlas (CGGA): A Comprehensive Resource with Functional Genomic Data from Chinese Glioma Patients (2021). Genomics, Proteomics & Bioinformatics 19(1):1–12.

[26] Zhang, K., Liu, X., Li, G,. Clinical management and survival outcomes of patients with different molecular subtypes of diffuse gliomas in China (2011– 2017): a multicenter retrospective study from CGGA. Cancer Biol Med 19, 1460-1476. 10.20892/j.issn.2095-3941.2022.0469. New

[27] Wang, Y., Qian, T., You, G., et al. (2015). Localizing seizure-susceptible brain regions associated with low-grade gliomas using voxel-based lesion-symptom mapping. NEURO-ONCOLOGY. 17(2): 282–288.

[28] Liu, X., Li, Y., Qian, Z., et al. (2018). A radiomic signature as a non-invasive predictor of progression-free survival in patients with lower-grade gliomas. NEUROIMAGE-CLINICAL. 20(1070-1077.

[29] Zhao, Z., Zhang, KN., Wang, QW., et al. Chinese Glioma Genome Atlas (CGGA): A Comprehensive Resource with Functional Genomic Data from Chinese GliomaPatients (2021). Genomics, Proteomics & Bioinformatics 19(1):1–12.

[30] Fang, S., Liang, J., Qian, T., et al. (2017). Anatomic Location of Tumor Predicts the Accuracy of Motor Function Localization in Diffuse Lower-Grade Gliomas Involving the Hand Knob Area. AMERICAN JOURNAL OF NEURORADIOLOGY. 38(10): 1990–1997

[31] Wang, Y., Wang, Y., Fan, X., et al. (2017). Putamen involvement and survival outcomes in patients with insular low-grade gliomas. JOURNAL OF NEUROSURGERY. 126(6): 1788–1794.

[32] Wishart, D. Metabolomics and the multi-omics view of cancer. Metabolites 2022, 12, 154.

[33] Ding, M.Q.; Chen, L.; Cooper, G.F.; Young, J.D.; Lu, X. Precision oncology beyond targeted therapy: Combining omics data with machine learning matches the majority of cancer cells to effective therapeutics. Mol. Cancer Res. 2018, 16, 269–278.

[34] Francescatto, M.; Chierici, M.; Rezvan Dezfooli, S.; Zandona, A.; Jurman, G.; Furlanello, C. Multi-omics integration for neuroblastoma clinical endpoint prediction. Biol. Direct. 2018, 13, 5.

[35] Lu, X.; Meng, J.; Zhou, Y.; Jiang, L.; Yan, F. Movics: An R Package for multiomics integration and visualization in cancer subtyping. Bioinformatics 2020, 36, 5539–5541.

[36] Rooney, M.S.; Shukla, S.A.; Wu, C.J.; Getz, G.; Hacohen, N. Molecular and genetic properties of tumors associated with local immune cytolytic activity. Cell 2015, 160, 48–61.

[37] Lacroix M, Abi-Said D, Fourney DR,et al. A multivariate analysis of 416patients with glioblastoma multiforme:prognosis, extent of resection, and survival. J Neurosurg. 2001;95:190–198.

[38] Lamborn KR, Chang SM, Prados MD. Prognostic factors for survival of patients with glioblastoma: recursive partitioning analysis. Neuro Oncol. 2004;6:227–235.

[39] Keime-Guibert F, Chinot O, Taillandier L, et al. Radiotherapy for glioblastoma in the elderly. N Engl J Med. 2007;356:1527–1535.

[40] Louis DN, Perry A, Wesseling P, et al. The 2021 WHO classification of tumors of the central nervous system: a summary. NeuroOncol 2021;23:1231–51.doi: 10.1093/neuonc/noab106 pmid: 34185076

[41] Hegi ME, Diserens AC, Gorlia T, et al. MGMT gene silencing and benefit from temozolomide in glioblastoma. N Engl J Med. 2005;352:997–1003.

[42] Eckel-Passow JE,Lachance DH,Molinaro AM, et al. Glioma groups based on 1p/19q, IDH, and TERT promoter mutations in tumors. N Engl J Med. 2015;372:2499–2508.

[43] Johnson DR, O’Neill BP. Glioblastoma survival in the United States before and during the temozolomide era. J Neurooncol. 2012;107:359–364.

[44] Koshy M, Villano JL, Dolecek TA, et al. Improved survival time trends for glioblastoma using the SEER 17 population-based registries. J Neurooncol. 2012;107:207–212.

[45] Taphoorn MJ, Sizoo EM, Bottomley A. Review on quality of life issues in patients with primary brain tumors. Oncologist. 2010;15:618–626.

